# Evolutionary insights into the emergence of virulent *Leptospira* spirochetes

**DOI:** 10.1101/2024.04.02.587687

**Authors:** Alexandre Giraud-Gatineau, Cecilia Nieves, Luke B. Harrison, Nadia Benaroudj, Frédéric J. Veyrier, Mathieu Picardeau

**Author notes:** corresponding author : Mathieu Picardeau, Institut Pasteur, « Biology of Spirochetes » unit, 28 rue du Dr Roux, 75714 cedex 15 Paris, France.

## Abstract

Pathogenic *Leptospira* are spirochete bacteria which cause leptospirosis, a re-emerging zoonotic disease of global importance. Here, we use a recently described lineage of environmental-adapted leptospires, which are evolutionarily the closest relatives of the highly virulent *Leptospira* species, to explore the key phenotypic traits and genetic determinants of *Leptospira* virulence. Through a comprehensive approach integrating phylogenomic comparisons with *in vitro* and *in vivo* phenotyping studies, we show that the evolution towards pathogenicity is associated with both a decrease of the ability to survive in the environment and the acquisition of strategies that enable successful host colonization. This includes the evasion of the human complement system and the adaptations to avoid activation of the innate immune cells. Moreover, our analysis reveals specific genetic determinants that have undergone positive selection during the course of evolution in *Leptospira*, contributing directly to virulence and host adaptation as demonstrated by gain-of-function and knock-down studies. Taken together, our findings define a new vision on *Leptospira* pathogenicity, identifying virulence attributes associated with clinically relevant species, and provide insights into the evolution and emergence of these life-threatening pathogens.

**AUTHOR SUMMARY:** *Leptospira* is a highly heterogeneous bacterial genus and leptospires are ubiquitous bacteria found as free-living saprophytes or as pathogens that can cause disseminated infections, from asymptomatic carriage in rats to lethal acute infection in both humans and animals. Leptospirosis is thus causing over one million cases and nearly 60,000 deaths annually. Despite leptospirosis being a re-emerging zoonosis, little is known about the ability of the etiologic agent to adapt to different hosts and cause disease. Here, combining genome analysis and phenotyping studies of representative species and mutant strains, we show that only a small group of species have the ability to evade the host immune system and cause disease. In addition, our findings provide key insight into the emergence of pathogens from a saprophytic ancestor through events of gene gain and genome reduction.

## INTRODUCTION

Spirochetes, which include the causative agents of Lyme disease, syphilis and leptospirosis, form an evolutionarily and morphologically unique phylum of bacteria. Despite their public health significance, spirochetes remain fastidious and challenging bacteria upon which to perform molecular genetic studies. As a result of their elusive nature, the underlying mechanisms for the emergence of these pathogens remain poorly understood.

Leptospirosis is a re-emergent zoonotic disease caused by pathogenic *Leptospira* species and accounts for approximately 1 million severe cases and 60,000 deaths every year [1]. The worldwide leptospirosis burden is expected to rise as climate and demographic changes fuel ideal conditions, including rising inequality which may contribute to rat-borne transmission which dominates human infection [1, 2]. It is likely that *Leptospira* infections are underdiagnosed due to the non-specific clinical presentations, poor performance of diagnostics tests and lack of notification systems and diagnostic laboratory capacity in most highly endemic countries [2]. Pathogenic leptospires typically infect humans via contact of abraded skin or mucous membranes with water contaminated by the urine of animal reservoirs, leading to significant morbidity in tropical and subtropical countries during rainy seasons and heavy rainfalls [3], as well as significant economic losses in the livestock industry [2]. Leptospirosis is associated with a wide spectrum of clinical manifestations, ranging from asymptomatic infection to a syndrome of multi-organ failure and death, and very little is known about the bacterial determinants implicated in severe infections [4].

The knowledge obtained from the study of model bacteria does not always translate to spirochetes. This is particularly true of pathogenic *Leptospira* spp. which are extracellular pathogens that lack many canonical virulence factors, including type 3 to type 10 secretion systems and associated effectors, pathogenicity islands, and virulence plasmids. Pathogenic *Leptospira* also have unique characteristics such as atypical lipopolysaccharides (LPS), peculiar defense mechanisms to oxidative stress, an endoflagellar system, and a large fraction of genes of unknown function [4–6]. Characterizing the mechanisms by which *Leptospira* evolved to be pathogenic should uncover novel mechanisms of bacterial virulence.

The last few years have seen several comparative genomics studies, revealing insights into the epidemiology, evolution, and genetic contents of *Leptospira* [7–11]. Notably, we have shown that the genus *Leptospira* is composed of 2 subclades (S1, S2) of free-living non-pathogenic species and 2 subclades (P1, P2) of species, including species with variable pathogenic potential (**Fig. 1a**). The subclade P1 can be further divided in two phylogenetically related groups termed P1+ (high-virulence pathogens) and P1- (low-virulence pathogens) [9]. Members of the P1+ group are established pathogenic species that have been reported to cause infections in both human and animals and the vast majority of *Leptospira* strains isolated from mammals belong to the P1+ group (**Fig. 1a**). Among the P1+, *L. interrogans* accounts for the majority of human infections and is, by far, the most studied of these organisms and consequently, most described virulence factors have been reported from this species. Interestingly, virulence-associated genes are over-represented in the entire P1 subclade with no clear distinction between the P1+ and the P1- groups (**Fig. 1a and Table S2**). Species in the P1- group and P2 subclade are mostly environmental isolates that have not been phenotypically well-characterized. Nevertheless, there is no evidence for an animal reservoir for most of the P1- and P2 species and only asymptomatic to mild infections have been sporadically reported [9, 11–16] (**Fig. 1a**). This defined phylogenetic structure, which mostly correlates with ecological niches and pathogenicity, provides a unique opportunity for the investigation of both the evolutionary events and the molecular mechanisms involved in the emergence of pathogenicity in *Leptospira*.

**Figure 1.**
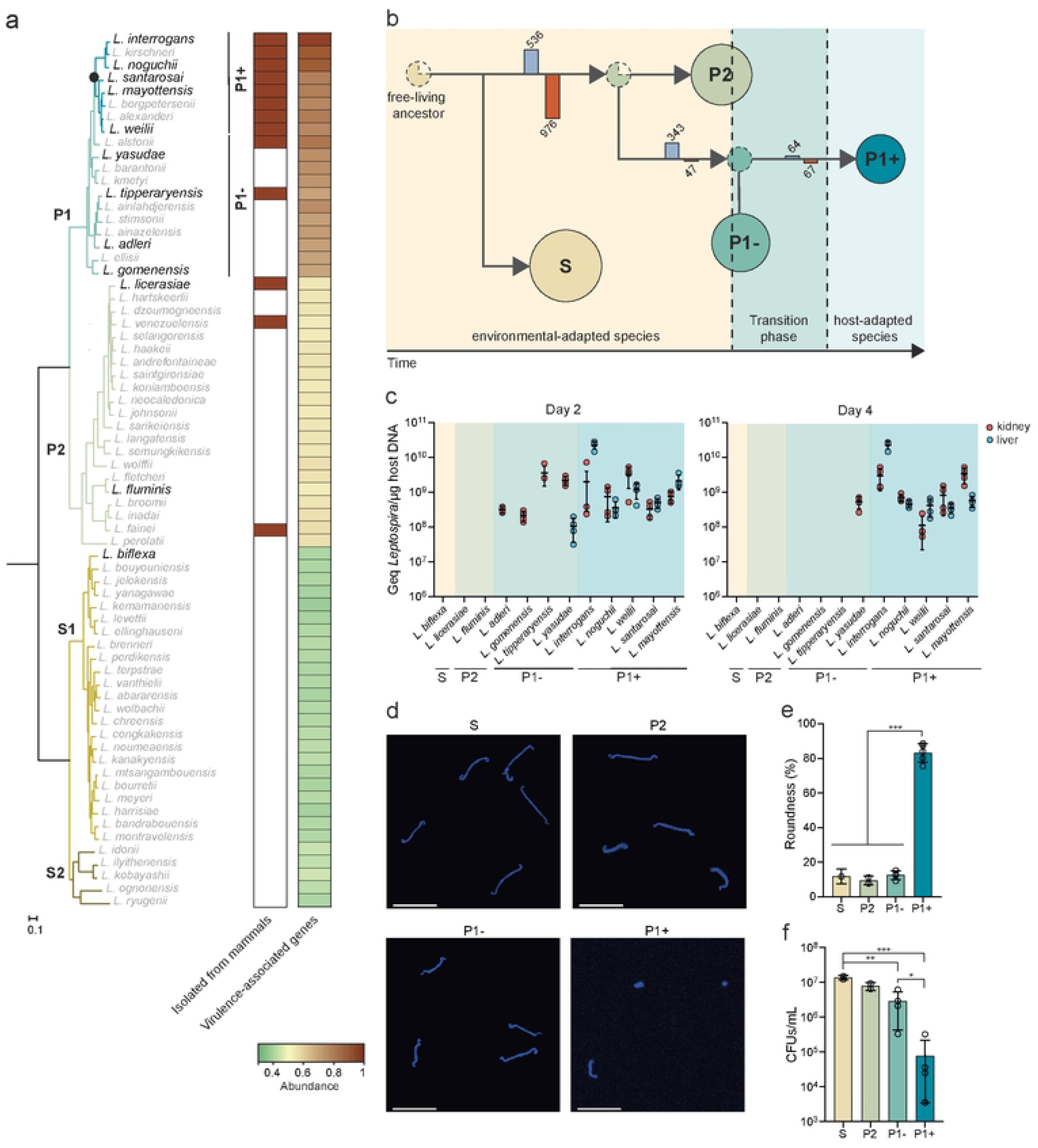
Phylogeny and evolution of host- and non-host adapted *Leptospira* lineages. **(a)** Phylogenetic tree based on soft-core genes (present in at least 95% of the genomes). The subclade P1, formerly referred to as the “pathogens” lineage, can be separated into two distinct groups: P1+ and P1-. P1+ consists of species associated with severe infections and diverged after a specific node of evolution (filled circle), while P1- comprises species that have not been isolated from patients and are considered as “low-virulent pathogens”. The species used in this study are indicated in the phylogenetic tree. Species in the P1- group and P2 subclade isolated from mammals are indicated by a red rectangle according to previous studies (*L*. *alstonii* (frogs) [18], *L. tipperaryensis* (shrew) [19], *L. licerasiae* (humans and rats) [20–22], *L. venezuelensis* (rodents, cattle and humans) [14], and *L. fainei* (pigs and wild boars) [23, 24]). Distribution of virulence-associated genes (**Table S2**) within the genus *Leptospira* are also shown using a heat map representation. **(b)** Evolutionary model and reconstruction of ancestral phenotypes in the genus *Leptospira* by PastML analysis using all maximum likelihood methods [25]. Branches are annotated with bars representing the sum of gene gain (blue bar) and loss (red bar). S, P2, P1- and P1+ clades and groups are indicated by spheres (whose size corresponds to the number of species) while most-recent common ancestors are indicated by dashed spheres. The dotted circles represent the most-recent common ancestors of each *Leptospira* group (S, P2, P1- and P1+), and the color indicates the most likely phenotype of that ancestor. **(c)** The virulence of *Leptospira* species was assessed by infecting hamsters (n = 4) with 10^8^ leptospires by intraperitoneal route. After 2 and 4 days of infection, the burden was assessed in kidney (red symbols) and liver (blue symbols) by quantitative PCR. Data are means ± SD; the absence of values indicates that *Leptospira* DNA was not detected. **(d-f)** Survival of *Leptospira* in water. Leptospires were incubated at RT in filter-sterilized spring water. At 21 days, leptospires were harvested, labelled with DAPI and analyzed by confocal microscopy **(d)** (scale bar: 10μm). The roundness of DAPI-positive leptospires was performed using Icy software **(e)** (n = 100 leptospires). The survival of *Leptospira* in filter-sterilized spring water after 21 days was determined by CFU **(f)**. S: *L. biflexa*; P2: *L. licerasiae*, *L. fluminis*; P1- group: *L. adleri*, *L. gomenensis*, *L. tipperyarensis*, *L. yasudae*; P1+ subgroup: *L. interrogans*, *L. noguchii*, *L. weilii*, *L. santarosai*, *L. mayottensis*.

The objectives of this study are to better characterize the P1- species, in particular with regard to their potential virulence, and to identify the genetic and phenotypic changes that characterize the emergence of P1+ species and their close associations with different hosts. This work provides an evolutionary framework for understanding the emergence of pathogenic *Leptospira* lineages.

## RESULTS

### Stepwise evolution of *Leptospira* from environmental saprophyte to life-threatening pathogen

To investigate the emergence of pathogens in *Leptospira*, we reconstructed the evolutionary history of the genus *Leptospira* and demonstrate that host-adapted pathogens evolved from environmental saprophytes (**Fig. 1b**). Our analysis shows that P1+ species emerged stepwise from clades of leptospires with P2 and subsequently P1- -like characteristics (**Fig. 1b**). Indeed, the most-recent common ancestor (MRCA) of the P (P1 and P2) clade is more likely to have P2-like phenotype than a P1- phenotype while the most recent common ancestor of the P1 clade is reconstructed to have a P1- phenotype.

To contrast their characteristics, we chose to study in more detail the phenotypes of 5 representative P1+ species (*L. interrogans, L. weilii, L. noguchii, L. mayottensis, and L. santarosai*) and 4 representative P1- species (*L. adleri, L. gomenensis, L. tipperaryensis, and L. yasudae*). Species sampled from P2 (*L. licerasiae and L. fluminis*) and S1 (*L. biflexa*) were also added in our analysis as additional reference strains (**Fig. 1 and Table 1**). We first assessed the burden of infection for representative species in the kidney and liver using the hamster model of acute leptospirosis. Although the P1- species can be detected at day 2 post-infection (pi), only the P1+ species were detected in organs at day 4 pi, with the exception of *L. yasudae* which was detected in kidneys only (**Fig. 1c**). S2 and P2 species are not detected at either time point. These results suggest that P1- species do not establish persistent infections in a susceptible host but do demonstrate a greater ability to survive in the host at very early times compared to P2 and S species.

**Table 1:** List of isolates used in this study.

| Species | Serogroup | Strain | Subclade/<br>group | Host | Source | GenBank |
| --- | --- | --- | --- | --- | --- | --- |
| *L. interrogans* | Pyrogenes | UP-MMC-NIID-LP | P1+ | Human | Philippines | GCA_001047635.1 |
| *L. mayottensis* | Mini | 200901116 | P1+ | Human | Mayotte | GCA_000306675.3 |
| *L. noguchii* | Panama | 201102933 | P1+ | Human | Guadeloupe | GCA_000306255.2 |
| *L. santarosai* | Shermani | 56164 | P1+ | Human | USA | GCA_000313175.2 |
| *L. weilii* | Celledoni | 14535 | P1+ | Human | Laos | GCA_000243995.3 |
| *L. adleri* | unknown | 201601302 | P1- | Soil | Mayotte | GCA_002811845.1 |
| *L. gomenensis* | unknown | KG8-B22 | P1- | Soil | New Caledonia | GCA_004770155.1 |
| *L. tipperaryensis* | unknown | GWTS#1 | P1- | Shrew | Ireland | GCA_001729245.1 |
| *L. yasudae* | unknown | M12A | P1- | Water | Mayotte | GCA_004770515.1 |
| *L. fluminis* | unknown | SCS5 | P2 | Soil | Malaysia | GCA_004771275.1 |
| *L. licerasiae* | unknown | VAR010 | P2 | Human | Peru | GCA_000244755.3 |
| *L. biflexa* | Semarang | Patoc | S1 | Water | Italy | GCA_000017685.1 |

We then wondered whether P1+ and P1- have the same ability to persist in the environment. After 21 days in spring water, P1+ species exhibited a 100-fold decrease in survival compared to other species and adopted a compromised round shape, while others species retained their helical shapes (**Fig. 1d-f**). In addition, the P1+ species showed a slower *in vitro* growth and reduced *in vitro* microbial and metabolic activity (redox activity, fluorescein diacetate hydrolysis and ATP production) in comparison to P1-, P2 and S species (**Fig. S1a-d**). Interestingly, a gradual decrease in microbial activity and ATP production is observed across *Leptospira* groups on the evolutionary trajectory towards P1+. This is consistent with the already described ongoing process of genome decay of P1+ [7–9, 17] characterized by an overrepresentation of mobile elements and pseudogenes, with 33% of non-functional genes being linked to metabolic processes (**Fig. S1e-h and Table S3**).

### Only P1+ species have developed strategies to escape host immunity

Pathogenic *Leptospira*, like most other pathogenic spirochetes, are stealthy pathogens that can escape the recognition by the host innate immune system [26]. We thus asked if P1- species had evolved similar strategies to escape or modulate host immunity during infection. First, we tested resistance to the complement system by assessing the survival of P1+ and non-infectious or low-virulent (P1-, P2 and S1) isolates in presence of normal human serum. Although, the majority of the genes encoding the resistance to the complement system described in *L. interrogans* are present in P1 and P2 but not in S (**Fig. S2**), only P1+ isolates resist to the complement-mediated killing and have the highest survival (86.27%±23.2) in human serum in comparison to other isolates (0.16%±0.01, 0.40%±0.28 and 1.83%±2.75 for S, P2 and P1-, respectively) (**Fig. 2a**). Analysis of Membrane Attack Complex (MAC) deposition on the cell membrane of *Leptospira* showed higher levels of deposition for S1, P2 and P1- compared to P1+ (83.00%±2.01, 81.25%±1.2, 65.64%±1.19 and 13.91%±1.21 for S, P2, P1- and P1+ respectively), which is associated with survival (**Fig. 2b**). Of note, the level of deposition for P1- is intermediate between P2 and P1+. Concordant with these results, maximal C3b deposition and bacterial opsonization in human macrophages was obtained in S1, P2 and P1- species (**Fig. S3a-b**).

**Figure 2.**
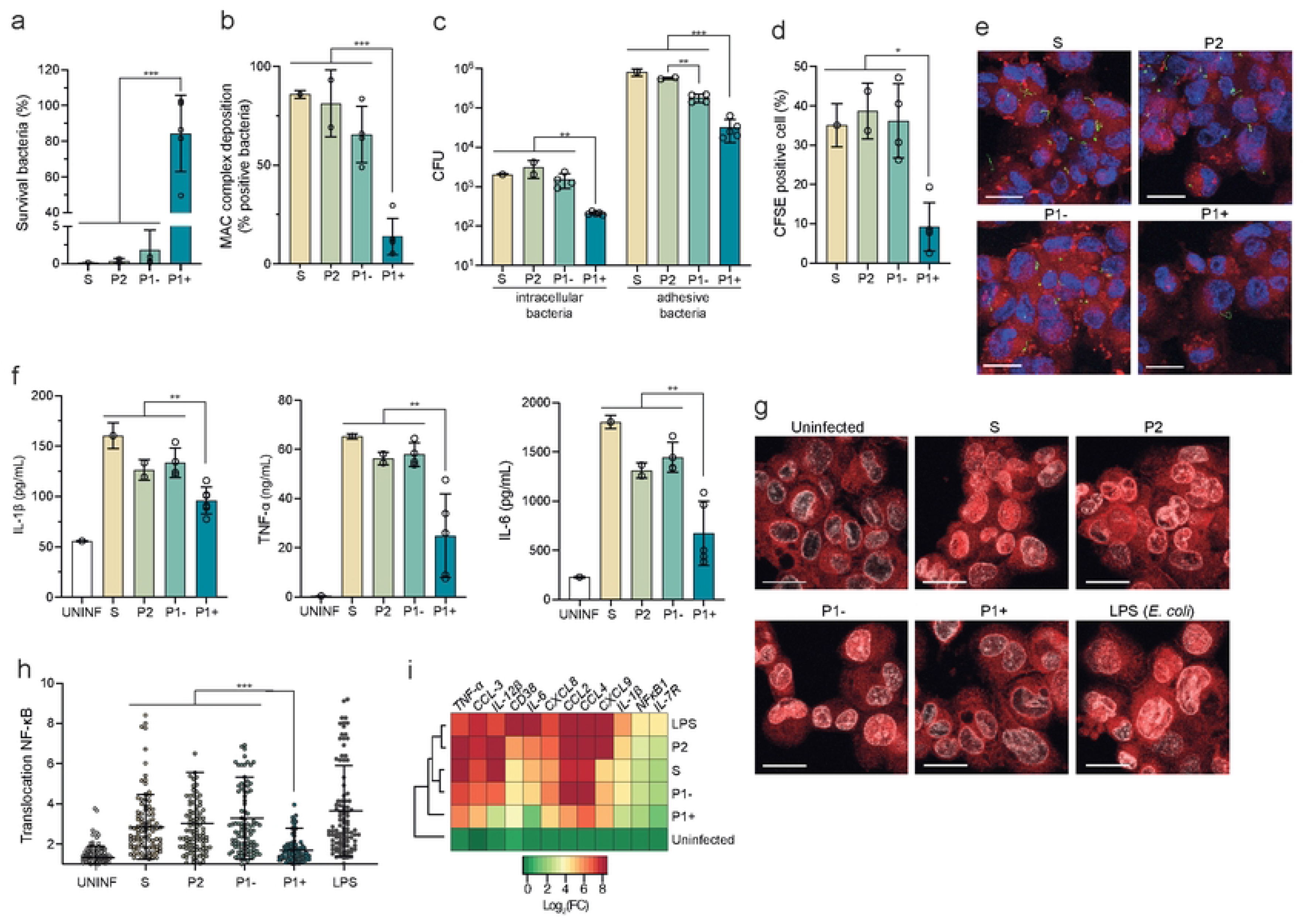
Only P1+ species escape the human complement system and have reduced interaction with human macrophages. **(a)** Survival of *Leptospira* upon exposure to human serum. Each *Leptospira* species was incubated in 20% of normal or heat-inactivated human serum for 2 hr. Lived bacteria were enumerated by CFU (counted in triplicate). The survival was compared to the inactivated-human serum. Unpaired two-tailed Student’s t test was used. ***p<0.0001. **(b)** MAC deposition in *Leptospira* was detected by indirect immunofluorescence. CFSE*-*stained *Leptospira* species were incubated with human serum for 30 min, fixed and then incubated with an anti-MAC antibody (C5b9). Indirect immunofluorescence was quantified by flow cytometry. The percentage was calculated by comparing the number of positive MAC-*Leptospira* to the number of negative MAC-*Leptospira*. Unpaired two-tailed Student’s t test was used. ***p<0.0001. **(c)** To assess bacterial internalization and adhesive bacteria, infected cells at 2 hr post-infection (PI) were washed with PBS and lysed directly before (adhesive bacteria) and after gentamicin treatment (bacterial internalization). Bacteria were enumerated by CFU (counted in triplicate). **(d-e)** Cells were infected with CFSE labelling leptospires. After 2 hr pi, cells were labelled with LysoTracker (Red). The fluorescence was analyzed by confocal microscopy. DAPI (blue) was used to visualize nuclei, CFSE (green) was used to visualize leptospires (scale bar: 20 μm). Quantification of CFSE positive macrophages compared to CFSE negative macrophages was performed using Icy software. Unpaired two-tailed Student’s t test was used. *p<0.01. **(f)** After 6 hr pi, IL-1β, TNF-α and IL-6 cytokines release from supernatant were measured by ELISA. Unpaired two-tailed Student’s t test was used. **p<0.001. **(g)** Representative fluorescence microscopy images of macrophages uninfected or infected with leptospires for 6 hr. Cells were labeled with antibody against NF-κB (red). DAPI (blue) and CFSE (green) were used to visualize nuclei and leptospires, respectively (scale bar: 20 μm). **(h)** Ratio between nuclear and cytosolic NF-κB fluorescence intensity (n > 100 cells per condition, two-way ANOVA test; ***p<0,01). LPS: *Escherichia coli* LPS **(i)** Heatmap showing relative expression of several genes regulated by NF-κB after 6hr pi for *Leptospira* infected cells. Expression of genes was analyzed and normalized using *gapdh* gene. Hierarchical clustering was performed using Ward’s method. Data are the mean ± SD (panels a-d, f, h, and i) or representative (panels e and g) of three independent biological replicates S: *L. biflexa*; P2: *L. licerasiae*, *L. fluminis*; P1- subgroup: *L. adleri*, *L. gomenensis*, *L. tipperyarensis*, *L. yasudae*; P1+ subgroup: *L. interrogans*, *L. noguchii*, *L. weilii*, *L. santarosai*, *L. mayottensis*.

Next, we showed that P1+ strains were significantly less internalized and less adherent to THP-1 macrophages than other species, (**Fig. 2c-e**). Concordantly, levels of pro-inflammatory cytokines (IL-1β, TNF-α and IL-6) were significantly lower in P1+-infected macrophages in comparison to other isolates, including P1- (**Fig. 2f**). To understand the molecular mechanisms underpinning the macrophage response, we analyzed the activation of the master transcriptional regulator of pro- and anti-inflammatory host response, NF-κB. P1+-infected macrophages showed a low level of NF-κB translocation into the nuclei (**Fig. 2g-h**). In contrast, NF-κB was mainly localized in the nucleus of S, P2 and P1- isolates. This was correlated with a higher induction of transcription of several inflammatory genes in non-P1+ isolates (**Fig. 2i**).

Taken together, these data show that only P1+ isolates have the capacity to resist the complement system and to avoid internalization by macrophages, which is correlated with a less severe early inflammatory response. In contrast, P1-, as well as P2 and S1 species, are sensitive to the complement system and are recognized by macrophages, triggering an inflammatory response.

### P1+-specific genetic determinants of *in vivo* virulence

Comparative genomics was then used to identify genes and/or pathways linked to the pathogenicity, revealing that the P1+ group is characterized by the acquisition of 64 genes and the loss of 67 genes (**Fig. 1b**, **Fig. 3a, Fig. S4 and Table S4**). Protein-encoding genes exclusively present in P1+ species, and potentially involved in the adaptation to the host, include uncharacterized proteins (26%), lipoproteins (20%), transposases (8%), and the established virulence factors collagenase [27], sphingomyelinases [28], and virulence-modifying (VM) proteins (19%) [29] (**Fig.3a**). Among these 64 specific genes, we found seven genes under positive selection (d*N*/d*S* >1; p-value <0.05) including genes encoding the collagenase, lipoproteins, VM proteins and an uncharacterized protein (**Fig. 3b and Table S5**). To experimentally evaluate the role of these proteins in host adaptation, we selected three genes (*colA*/*LIMLP_03665, hyp/LIMLP_09380* and *VM/ LIMLP_11655*) that are induced during *in vivo*-like conditions (**Fig. S5**) as previously shown [30–32] (**Fig. S6).** Each gene was heterologously expressed in the P1- species *L. adleri* and *L. yasudae*. Expression was confirmed by RT-qPCR (**Table S6**) and *colA/LIMLP_03665-*expressing strains exhibited collagenase activity (**Fig. S7**). Production of the three *L. interrogans* proteins led to increased burdens of the low-virulent strains in hamsters in at least one of the tested conditions (**Fig. 3c-d**) whereas CRISPR-*dcas9*-based transcriptional silencing of *colA/LIMLP_03665*, *VM/LIMLP_11655* and *colA*/*LIMLP_03665* in the pathogen *L. interrogans* resulted in an attenuation of virulence in the hamster model (**Fig. S8 & Fig. 3h**). We also found that *hyp/LIMLP_09380* expression in P1- strains induced resistance to complement-mediated killing and lowered MAC deposition (**Fig. 3e & Fig. S9**), while *VM/ LIMLP_11655* expression reduced the inflammatory response of P1- strain-infected macrophages (**Fig. 3f-g**). The role of *hyp/LIMLP_09380* in inducing resistance to complement-mediated killing and the involvement of *VM/ LIMLP_11655* in reducing the inflammatory response of infected macrophages in the pathogen *L. interrogans* were validated with the respective knock-down mutants (**Fig. S10)**.

**Figure 3.**
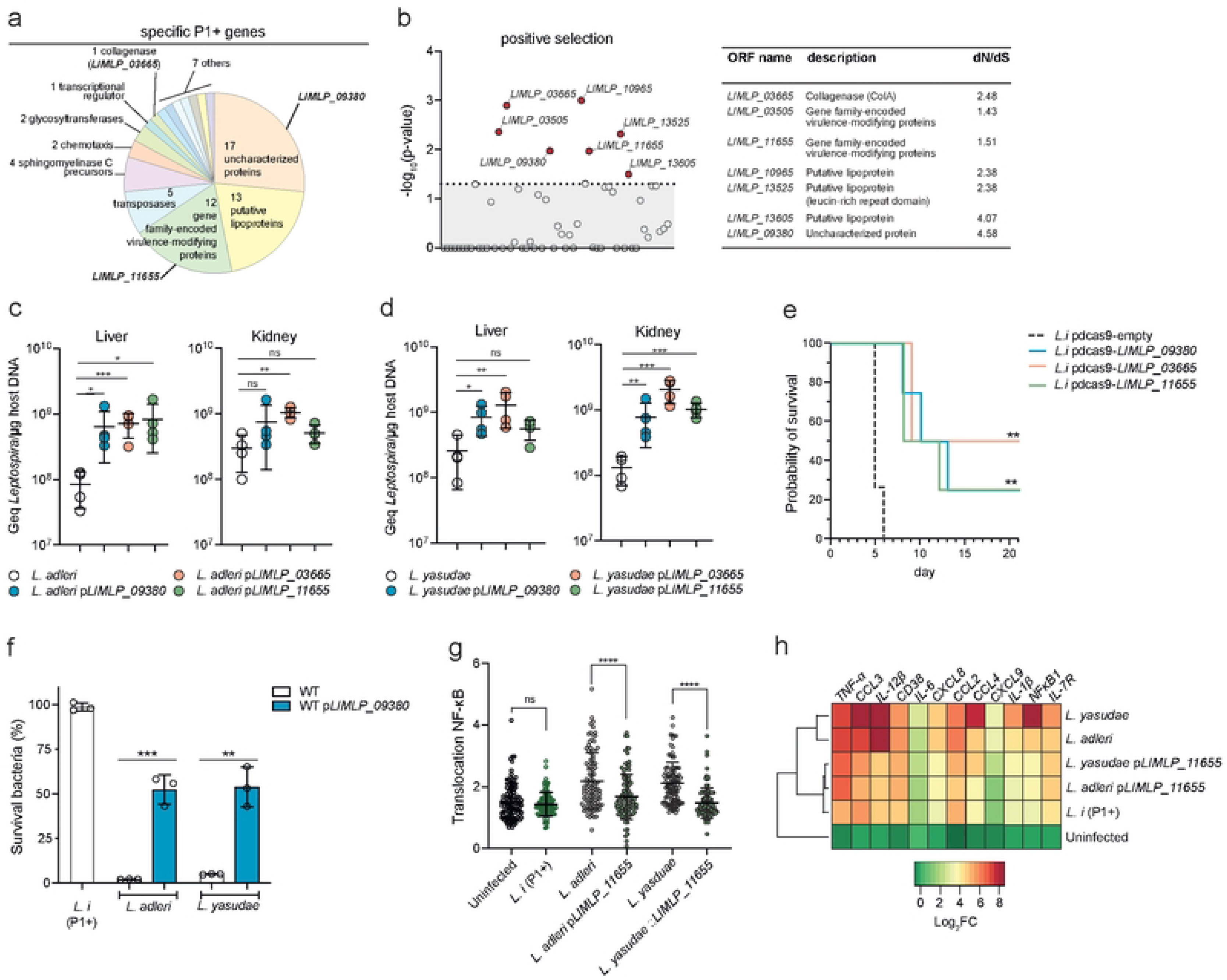
Acquisition of specific P1+ genes involved in host-associated lifestyle. **(a)** Distribution of the 64 P1+-specific genes **(b)** Positive selection analysis of the P1+-specific genes. Significative positive selection was determined using the PoSeiDon pipeline. Significant genes (p-value <0.05, dashed line) and rate of non-synonymous (d*N*) and synonymous (d*S*) in the alignment of orthologous sequences are indicated. **(c-d)** The virulence of *L. adleri* (c) and *L. yasudae* (d) P1- strains was assessed by infecting hamsters (n = 4) by intraperitoneal route with 10^8^ leptospires. After 1 day of infection, leptospiral load in kidney and liver was assessed by quantitative PCR. Unpaired two-tailed Student’s t test was used.*p< 0.01, **p<0.001, ***p<0.0001, ns: not significant. **(e)** Survival of hamsters (n = 4) infected intraperitoneally with 10^6^ *Leptospira* for each construct. Statistical significance in comparison with *L.i* pdcas9-empty was determined by a Log rank Mantel Cox test (**p<0.0021). **(f)** Effect of hyp/LIMLP_09380 on survival in human serum. P1- species *(L. adleri* and *L. yasudae*) producing or not *LIMLP_09380* were incubated in 20% of human serum or inactivated-human serum for 2 hr; *L. interrogans* WT (*L.i*) is shown here as a reference for P1+ species. After incubation, the bacteria were enumerated by CFU (counted in triplicate). The percentage of surviving bacteria was calculated using the inactivated-human serum as normalization. Unpaired two-tailed Student’s t test was used. **p<0.001, ***p<0.0001. **(g-h)** Effect of VM/LIMLP_11655 on innate immune response of human macrophages. Ratio between nuclear and cytosolic NF-κB fluorescence intensity (n > 100 cells per condition, two-way ANOVA test; ****p<0,001; ns: not significant) in the different *Leptospira* strains (f). Heatmap showing relative expression of several genes regulated by NF-κB after 6hr pi for *Leptospira* infected cells (g). Expression of genes were analyzed and normalized using *gapdh* gene. Hierarchical clustering procedure of *Leptospira* genus was performed using Ward’s method. **(h)** Survival of hamsters (n = 4) infected intraperitoneally with10^6^ *Leptospira*.

## DISCUSSION

Pathogenic *Leptospira* (P1+) evolved from environmental bacteria in progressive trajectory of host-adaptation to animals through deep-time, likely beginning with the appearance of the first mammals [3]. Other *Leptospira* species are mostly environmental isolates but some P2 and P1- species may be responsible for asymptomatic to mild infections in both humans and animals [12–16, 21, 22]. Our results, when summarized and subjected to a multivariate hierarchical clustering analysis, highlight a distinct separation of P1+ isolates from others *Leptospira* subgroups. Interestingly, the P1- isolates, which possess most of the known virulence factors and are phylogenetically closely related to P1+, were clustered with the non-infectious or low-virulent P2 and S isolates (**Fig. 4**).

**Figure 4.**
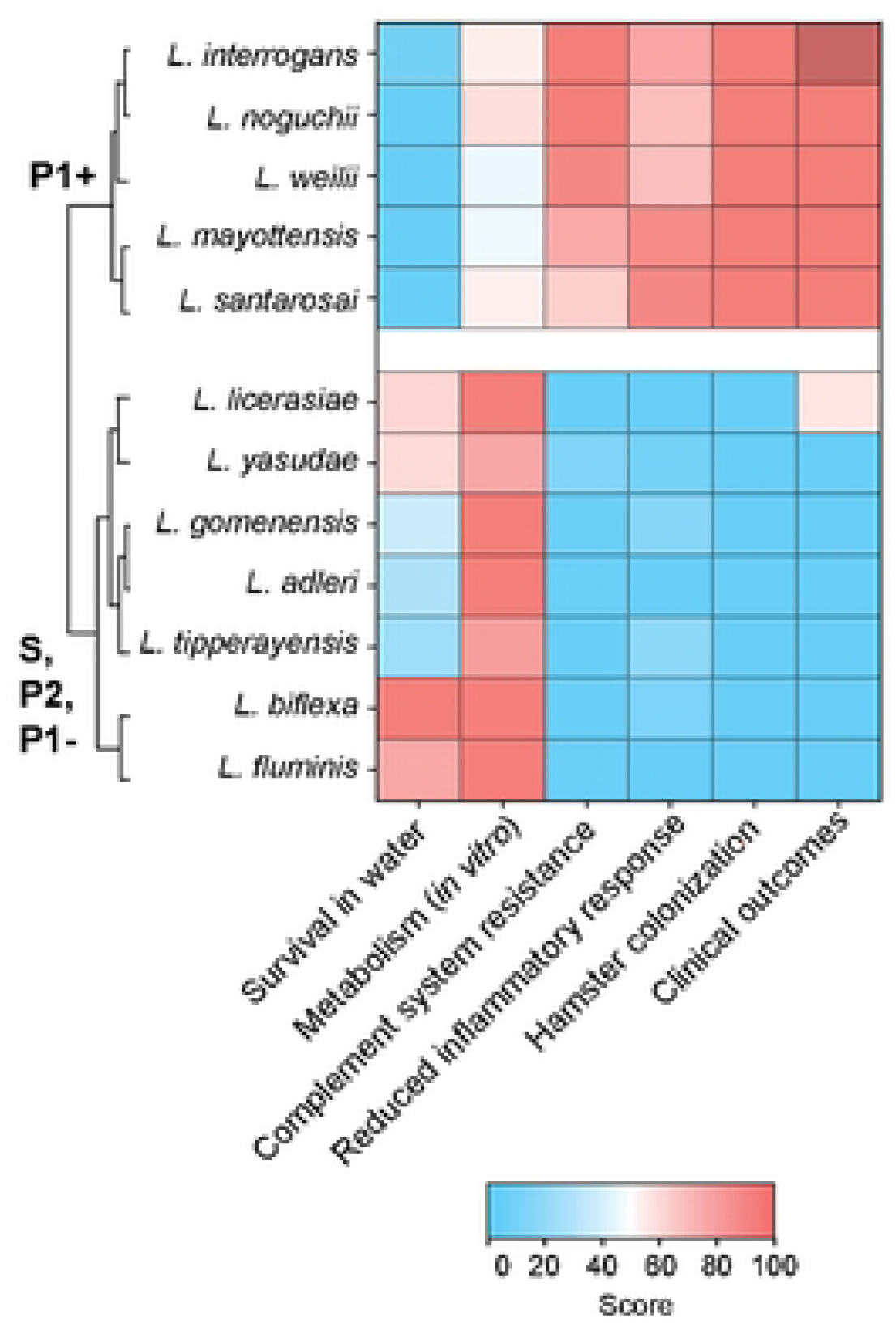
Heatmap representation of the main features of representative *Leptospira* species described in this study. With the exception of *L. licerasiae* [20–22], only P1+ species are responsible for infections in humans. Hierarchical clustering was performed using Ward’s method.

We found that only P1+ strains can establish persistent colonization in the acute animal model of infection showing that these bacteria possess mechanisms enabling the spirochete to survive longer in the blood and to proliferate in target organs, consistent with previous observations [11]. Along the same lines, we show that only the P1+ species can escape immune surveillance and complement-mediated killing. Structural differences within the LPS lipid A may contribute to differential recognition by host immune cells observed between P1+ and P1-/P2 species [33] (**Fig.5**). Adaptation of P1+ species to a wide range of hosts, unlike other species, also correlates with a more diverse and complex LPS O-antigen biosynthesis gene locus [7].

**Figure 5.**
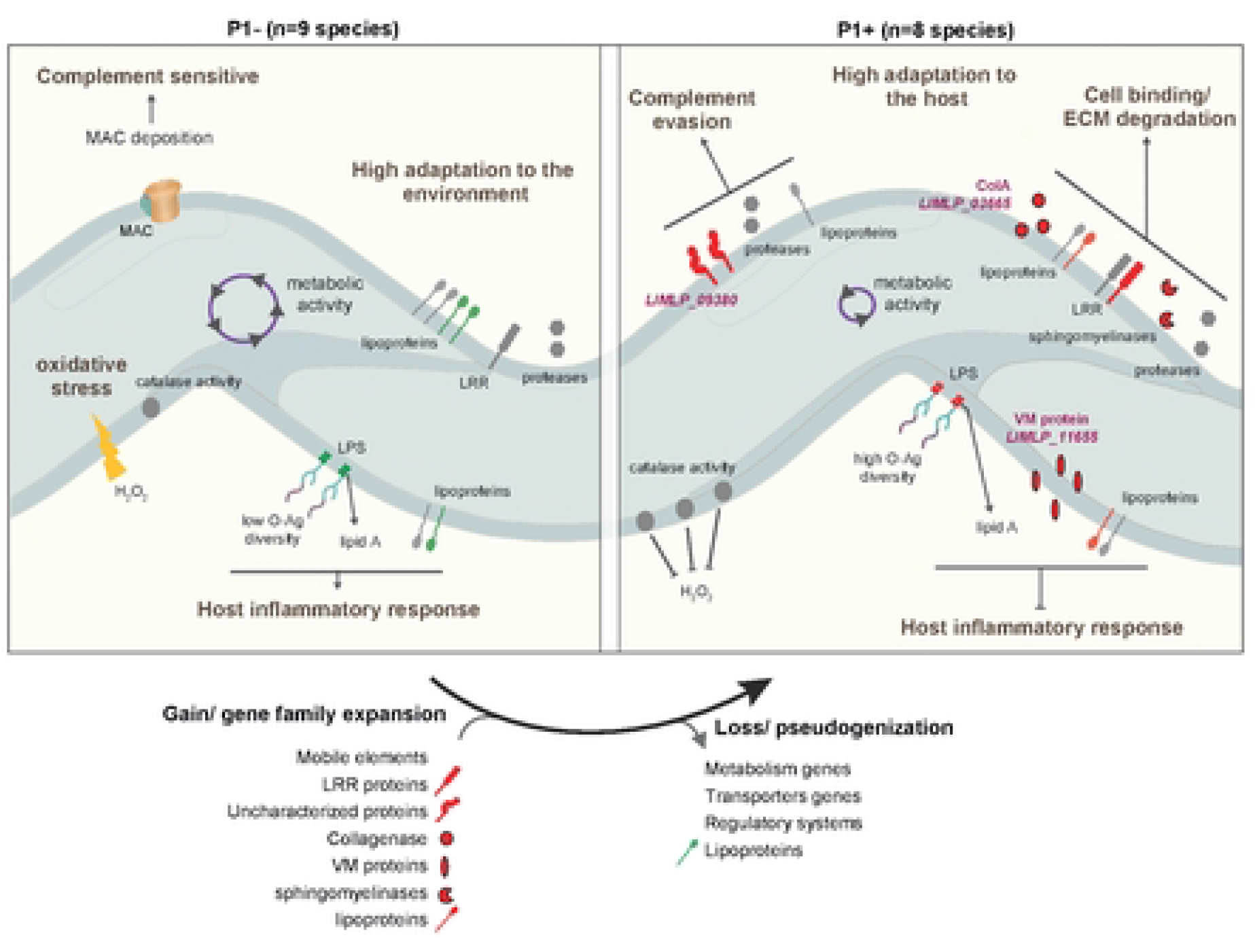
Schematic representation of the evolutionary transition from environmental-adapted *Leptospira* species (P1- group) to host-adapted *Leptospira* species (P1+ group). Host-associated factors found in both P1- and P1+ species (Fig. S2) are indicated in grey. Factors found exclusively in P1+ and P1- are indicated in red and green, respectively. Specific lipoproteins, Leucin-rich repeat (LRR)-encoding proteins, sphingomyelinase-like proteins, virulence-modifying (VM) proteins and uncharacterized proteins are prominent among P1+ isolates. The different factors contributing to host adaptation of P1+ species are represented, including a collagenase (encoded by LIMLP_03665), a hypothetical protein (encoded by LIMLP_09380) and a VM protein (encoded by LIMLP_11655). LIMLP_09380 participates in evasion from complement-mediated killing and the VM proteins is involved in prevention of host inflammatory response. In addition, several factors are contributing in the cell binding and ECM (extracellular matrix) degradation for P1+ species. The lipopolysaccharides (LPS) of P1+ species, which have a higher complexity than those of other *Leptospira* species [7, 33] may differentially interfere with the host and confer escaping from immune surveillance. The reduced *in vitro* microbial and metabolic activities of P1+ species in comparison to P1- species might be also important for adaptation to the host. The catalase activity of P1+ species is higher than in P1- species, allowing them to better tolerate H_2_O_2_ as encountered inside a host.

Although P1-, P2 and S species are cleared by host defenses at day 4 pi, P1- species can be differentiated from P2 and S1 species as they can be detected in the early stage of infection, suggesting a limited degree of host adaptation or pre-adaptation in response to other uncharacterized selective pressures. Thus, it has been shown that adaption of environmental microbes to a host-independent factor can incidentally increase their ability to cause an infection [34].

Our findings suggest that the gradual evolution from P1-like species to P1+ with the capacity for host colonization was based on only a few key genetic evolutionary innovations through the loss and/or the acquisition of genes, along with gene family expansion and pseudogenization. Putative virulence factors specific to P1+ isolates include families of sphingomyelinase-like proteins and leucine-rich repeat (LRR) proteins, both probably mediating host-pathogen interactions [28, 35]. We also show experimentally that, among the specific genetic determinants of P1+, at least three genes (*hyp/LIMLP_09380, colA/LIMLP_03665,* and *VM/LIMLP_11655)* actively contribute to *in vivo* virulence through evasion of host defense. They include a protein belonging to a paralogous family of proteins called Virulence-Modifying proteins (VM proteins) which are not present in P1- and expanded in the most virulent species (P1+) whose genomes can encode 12 or more paralogs [29], as well as a collagenase which, when mutated, reduces virulence [27]. On the other hand, genes lost in P1+ isolates are predominantly associated with metabolism (31%, **Fig. S4 and Table S4**) and this is corroborated with reduced metabolic activity and ability to survive in water when compared to other *Leptospira* species. At one extreme of this evolutionary trajectory is the obligate pathogen *L. borgpetersenii*, which cannot persist in the environment and whose genome is characterized by gene decay through extensive IS-mediated genome rearrangement and pseudogenization [17]. Most P1+ species, howsoever, can persist in the environment and represent different intermediate stages of genome reduction relative to *L. borgpetersenii* on the evolution transition to an obligate parasitic lifestyle.

Genome evolution is a key determinant in host adaptation, but other mechanisms should also be considered. Previous studies had demonstrated that genes controlling adhesion, uptake and intracellular survival within macrophages and neutralization of the complement system [36, 37] are involved in *Leptospira* pathogenicity. However, we have shown here that most of these genes involved in virulence and host adaptation are also present within the P1- group (**Fig. S2**), implying that differential gene expression of some virulence determinants might also be at the origin of the different pathogenicity of P1- and P1+ species. Supporting this hypothesis, we have recently demonstrated that P1+ isolates exhibit greater resistance to peroxide, a characteristic associated with a constitutive higher expression of *katE*-encoding catalase compared to P1- isolates, which display a lower tolerance to peroxide [38]. Therefore, a rewiring of transcriptional circuits could also contribute to the emergence of pathogenicity in *Leptospira* genus.

In summary, our study shows that the evolution of virulence in leptospires, defined here as the ability to adapt and persist in the host, occurred with the successive appearance of intermediate phenotypes (**Fig. 5**). Firstly, a reduction in the metabolism-related genetic repertoire occurred in the most-recent common ancestor of P2 species persisting through the phylogenetic grade of P1- species until the emergence of P1+ species. Secondly, mechanisms that support host colonization evolved in the most-recent common ancestor of the P1 clade, as demonstrated by the detection of P1- species in the organs of the acute animal model during the early stages of infection. Finally, an enrichment in mobile elements appears in P1+ species, which contributes to reductive genomic evolution. This phenomenon further reduces the content of metabolic genes and marks a crucial stage in the evolution of *Leptospira*, as these species adapt to different hosts by acquiring genes essential for evading the host immune response and experience a concomitant relaxation of purifying selection on genes important for survival in the environment. Of note, our results suggest that P1- species could, under certain conditions (e.g. immunocompromised individuals), be responsible for infections in humans. However, the diagnostic tools currently available are not capable of detecting P1-/P2 isolates, and therefore their relevance to human and animal health has yet to be determined. Similarly, the sporadic reports of P1-/P2 infection in animals need to be further investigated to determine whether some animals may serve as reservoirs for low-virulent species. In addition, all P1+ species are probably not equally virulent. Further studies should include large scale analyses integrating clinical data, phenotypic analysis and comparative genomics of P1+ isolates to provide a better understanding of bacterial factors associated with severe infections.

Our findings refine our understanding of virulence in *Leptospira* and provide novel insights into the pivotal steps of host adaptation in *Leptospira*. This study, which enables us to better distinguish hypervirulent *Leptospira* strains from others, may also have important public health implications.

## MATERIAL and METHODS

### Bacterial strains and culture conditions

*Leptospira* strains used in this study are indicated in Table 1. *Leptospira* strains were cultivated aerobically in Ellinghausen-McCullough-Johnson-Harris liquid medium (EMJH) at 30°C with shaking at 100 rpm or onto 1% agar solid EMJH media at 30°C.

### Survival in water

Exponentially growing *Leptospira* species were centrifuged at 2,600 g for 15 min, washed three times and resuspended into filter-sterilized spring water (Volvic). All *Leptospira* species were adjusted to 5×10^8^ leptospires/ml and incubated at room temperature (RT) in the dark. At 21 days, survival was determined by enumeration of colony-forming unit (CFU) on EMJH agar plates. For staining, bacteria were fixed with 4% paraformaldehyde at RT for 15 min, stained with DAPI (1µg/ml, Thermo Fisher) for 10 min and mounted on a slide using Fluoromount mounting medium (Thermo Fisher). Quantification of DAPI staining and roundness was performed using Icy software.

### In vivo animal studies

Four week-old Syrian Golden hamsters (RjHan:AURA, Janvier Labs) were infected (4 per group) by intraperitoneal injection with 10^6^ or 10^8^ *Leptospira* as enumerated using a Petroff-Hausser counting chamber. The animals were monitored daily and euthanized by carbon dioxide inhalation upon reaching the predefined endpoint criteria (sign of distress and morbidity). To assess leptospiral load, blood, kidney, and liver were sampled and DNA was extracted with the Tissue or Blood DNA purification kit (Maxwell, Promega). The bacterial burden and host DNA concentration were determined by qPCR with the Sso Fast EvaGreen Supermix assay (Bio-Rad) using the *flaB2* and *gapdh* genes, respectively. *Leptospira* load was expressed as genomic equivalent (GEq) per µg of host DNA.

### Metabolism activity and enzymatic assays

Exponentially growing *Leptospira* (2×10^8^) were incubated in EMJH at 30°C. Rezasurin (Alamar Blue Assay, ThermoFisher) was added, and bacteria were incubated for 8 hr. The absorbance was measured at 570 and 600 nm. Redox activity was determined based on the ability of cells to reduce rezasurin into resorufin following the manufacturer’s instructions.

Exponentially growing *Leptospira* (2×10^9^) were incubated with fluorescein diacetate (Sigma-Aldrich) at 2 mg/ml in acetone at 30°C for 10 min, then a 2:1 chloroform/methanol solution was added. After centrifugation at 5,000 g for 5 min, the aqueous phase was recovered, and the microbial activity (esterase activity) was obtained by measuring the absorbance at 490nm.

Exponentially growing *Leptospira* (2×10^9^) were used to determine ATP concentration using the luminescent ATP detection assay (Abcam), according to the manufacturer’s instructions.

Total extracts or the supernatant of *Leptospira* (5.5 µg) was used to determine the collagenase activity using the Collagenase Activity Assay Kit (Abcam), according to the manufacturer’s instructions.

### Comparative genomics and phylogeny

Gene acquisition and loss were assessed using an in-house developed software, MycoHIT [39]. Tblastn searches (E-value = 1e^-10^) were independently performed for all 68 *Leptospira* species against reference protein-coding sequences of i) *L. interrogans* str. 56601 (to evaluate gene acquisition), and ii) *L. biflexa* str. Patoc 1 (Paris) (for assessment of gene loss). Presence/absence of genes was determined by a 60% similarity threshold, observed in at least 80% of the members within the target group. Furthermore, to ensure specificity, presence/absence of these genes should not surpass 20% in other groups. To illustrate, for a gene to be designated as “present” in the P1+ group, it needed a similarity ≥ 60% in a minimum of 7 out of 8 species within that group.

Additionally, the presence of that gene could not exceed 20% prevalence across the species included in P1-, P2, S1, and S2 (i.e., present in < 12 species).

Protein-coding genes within each species were classified based on the Clusters of Orthologous Genes (COG) database using eggNOG mapper (options: --evalue 0.001 --score 60 --pident 40 -- query_cover 20 --subject_cover 20 --target_orthologs all) [40]. The representativeness of COG categories per species was calculated as the ratio of protein-coding genes within each specific category, normalized by the total number of protein-coding genes in the respective species.

Pseudogene prediction was performed using Pseudofinder version 1.1.0 (option --annotate) [41]. The search procedure involved sequence alignment against the UniProt/TrEMBL protein database through DIAMOND (option --diamond within the Pseudofinder command line) [42]. Functional characterization of identified pseudogenes was performed based on COG classification, as previously described.

Representative genomes of *Leptospira* species are indicated in Table 1. The gene orthology was determined using GET_HOMOLOGUES version 2021 using parameters of minimum protein coverage of ≥ 70%, E-value = 0.01.

Protein motifs were detected using InterProscan v5.30_69.0 (https://doi.org/10.1093/bioinformatics/btu031) for each genomic reference of the 68 *Leptospira* species.

Ancestral reconstruction of the phenotypes of the *Leptospira* genus was performed using PastML [25]. First, a core genome alignment of 68 species of the *Leptospira* genus, encompassing members from P1+, P1-, P2, S1 and S2, was performed using MAFFT through Roary v3.11.2. [43]. The alignment parameters included a 60% identity cut-off and required gene prevalence in at least 95% of the analyzed genomes (Roary’s options: -e –mafft -I 60 -cd 95). The resulting alignment comprised a total of 624 soft-core genes, which was used for subsequent phylogenetic analysis. The best evolutionary model was determined using ModelFinder within IQ-TREE version 1.6.11 [44]. The best-fit model was identified as GTR + F + R7 by Bayesian Information Criterion (BIC).. Maximum-likelihood phylogenetic analysis was performed with IQ-TREE, incorporating 10,000 ultrafast bootstrap replicates [45]. The final phylogenetic tree was visualized with FigTree v1.4.4 (http://tree.bio.ed.ac.uk/software/figtree/) and rooted with the saprophytic group (S, comprising S1 and S2as shown in Vincent et al. [9]). This rooted phylogenetic tree, along with a csv file associating species with taxonomic groups (P1+, P1-, P2, S1 and S2), served as inputs for PastML. Ancestry prediction was performed by using a combination of all maximum-likelihood methods (maximum a posteriori, MAP; joint; marginal posterior probabilities approximation, MPPA), and the F81 model.

### Positive selection of P1+ specific genes

dN/dS was calculated using codeML though the PoSeiDON pipeline (10.1093/bioinformatics/btaa695). This pipeline performed in-frame alignment of each protein-coding sequence, phylogenetic reconstructions and detection of positively selected sites in the full alignment. Maximum-likelihood tests to detect positive selection under varying site models are performed using M7 versus M8 by codeML with three independent codon models F1X4, F3X4 and F6. Then, we used an empirical Bayes approach to calculate posterior probabilities that a codon coming from a site class with dN/dS>1. Genes were considered to be positively selected when the p-value is <0.05 and the dN/dS ratio exceeds one.

### Complement system activity

*Leptospira* strains were incubated with 20% human serum from healthy individual donors or 20% heat-inactivated human serum diluted in PBS. At the indicated times, bacteria were harvested by centrifugation at 2,600 g for 15 min and fixed with 4% paraformaldehyde for 15 min at RT. Bacteria were incubated with anti-C5b9 (Thermo Fisher) or with anti-C3b (Thermo Fisher) for 2 hr at RT. Bacteria were washed three times with PBS and incubated with Alexa Fluor 555 secondary antibody (Thermo Fisher) for 2 hr at RT. Fluorescence was analyzed using a CytoFLEX Flow Cytometer (Beckman Coulter). The analysis was performed using the FlowJo software.

### Macrophages infection model

THP-1 (ATCC® TIB-202) cells were cultured in RPMI 1640 medium (Gibco) supplemented with 10% heat-inactivated fetal bovine serum (Sigma) and 2 mM L glutamine (Gibco). For all experiments, THP-1 cells were differentiated/activated into macrophages by a treatment with 50 nM PMA for 2 days following by a 24 hr incubation without PMA. Activated macrophages were inoculated with *Leptospira* at a multiplicity of infection (MOI) of 100 bacteria-per-cell during 2 hr following by 1 hr of gentamicin treatment (Sigma) at 100 µg/ml. Before and after gentamicin treatment, cells were washed three times with PBS.

For Carboxyfluorescein succinimidyl ester (CFSE) labelling, leptospires were resuspended in PBS with 5 µM CFSE (Sigma-Aldrich) for 30 min at RT and then washed three times in PBS. After infection, cells were incubated with 100 nM LysoTracker DND-99 (Thermo Fisher) for 1 hr at 37°C. THP-1 cells were washed three times with PBS and then fixed with 4% paraformaldehyde at room temperature for 15 min. Nuclei were stained with DAPI (1 µg/ml, Thermo Fisher) during 10 min and mounted on a side using Fluoromount mounting medium (Thermo Fisher). Fluorescence was analyzed using a Leica TCS SP8 Confocal System. Quantification of CFSE-positive cell was performed using Icy software.

### Host immune response

Quantification of cytokines present in cell culture supernatants was performed by ELISA using the DuoSet ELISA kit (R&D Systems) for TNF-α, IL-1β and IL-6, according to the manufacturer’s instructions.

Analysis of NF-KB activation was performed at 6 hr post-infection. Cells were fixed with 4% paraformaldehyde at RT for 15 min, incubated for 5 min with 0.5% saponin (Sigma-Aldrich) in PBS and then incubated for 30 min in 1% BSA (Sigma-Aldrich) and 0.1% saponin (Sigma-Aldrich) in PBS, to permeabilize the cells and to block nonspecific binding. Cells were incubated with anti-NF-kB antibody (Thermo Fisher) overnight at 4°C. Cells were washed and incubated with Alexa Fluor 555 secondary antibody (Thermo Fisher) for 2 hr. Nuclei were stained with DAPI (1 µg/ml) for 10 min. After labeling, coverslips were set in Fluoromount G medium (Thermo Fisher). Fluorescence was analyzed using a Leica TCS SP8 Confocal System and the NF-kB translocation analysis was performed using Icy software.

### Measure of gene expression

Total RNAs from *Leptospira* species or macrophages were extracted using QIAzol lysis reagent (Qiagen) and purified with RNeasy columns (Qiagen). Reverse transcription of mRNA to cDNA was carried out using the iScript™ cDNA Synthesis kit (Bio-Rad), followed by cDNA amplification using the SsoFast™ EvaGreen® Supermix (Bio-Rad). All primers used in this study are listed in Supplementary Table 1. Reactions were performed using the CFX96 real-time PCR detection system (Bio-Rad). The relative gene expression was assessed according to the 2^-ΔCt^ method using *flaB2* (*LIMLP_09410*) or 16S RNA (*LIMLP_04870*) as reference genes for *Leptospira* or *gadph* as reference gene for human macrophages.

### Genetic manipulations of *Leptospira*

Heterologous expression of *LIMLP_09380*, *LIMLP_03665* and *LIMLP_11655 L. interrogans* genes in *L. adleri* and *L. yasudae* was performed by cloning the genes in pMaGRO [46] and introducing the conjugative plasmids in *Leptospira* as previously described [47]. *Leptospira* conjugants were selected on EMJH agar plates containing 50 µg/ml spectinomycin.

Gene silencing in *L. interrogans* was performed using the leptospiral replicative vector pMaOri.dCas9 [48]. The sgRNA cassettes targeting *LIMLP_09380 (5’TGTGGTACTATGAGATATAC3’)*, *LIMLP_03665 (5’TCGGTTATGACTTTCGGACC3’)* and *LIMLP_11655 (5’TTTGGACTGCTGACGGAAAG3’)* were cloned into the replicative plasmid pMaori.dcas9 and plasmids pMaori.dcas9_sgRNA-*LIMLP_09380*, pMaori.dcas9_sgRNA-*LIMLP_03665* and pMaori.dCas9_sgRNA-*LIMLP_11655* were introduced into *L. interrogans* by conjugation. *Leptospira* conjugants were selected on EMJH agar plates containing 50 µg/ml spectinomycin.

### Quantification and statistical analysis

Data are expressed as means ± standard deviations (SD). Statistical analyses were performed with Prism software (GraphPad Software Inc.), using the t test and one-way analysis of variance (ANOVA) as indicated in the figure legends.

### Ethics Statement

Sera were obtained from healthy donors. The blood collection was carried out in accordance with the approved French Ministry of Research and French Ethics Committee protocols by the Etablissement Français du Sang (EFS, n°18/EFS/041). Protocols for animal experiments are conformed to the guidelines of the Animal Care and Use Committees of the Institut Pasteur (Comité d’éthique d’expérimentation animale CETEA # 220016), agreed by the French Ministry of Agriculture. All animal procedures carried out in our study were performed in accordance with the European Union legislation for the protection of animals used for scientific purposes (Directive 2010/63/EU).

## ACKNOWLEDGEMENTS

We thank Jean François Mariet for controlling human sera. AGG was funded by a PTR2019-310 grant (NB). This research was supported by the Institut Pasteur through grant PTR 30-2017 (MP and FJV), the National Institutes of Health grant P01 AI 168148 (MP and FJV) and the French National Research Agency, grant number SPIraL-19-CE35-0006-01. CN received a Ph.D. studentship Calmette & Yersin from the Institut Pasteur International Network. LBH is supported by the Fonds de Recherche du Québec-Santé: Clinician Scientist Training Program for Residents in Medical Specialties. FJV received a Junior 1 and Junior 2 research scholar salary award from the Fonds de Recherche du Québec-Santé. The funders had no role in study design, data collection and analysis, decision to publish, or preparation of the manuscript.

